# Plasma concentrations of nirmatrelvir and molnupiravir required for inhibition of SARS-CoV-2 replication differ between rhesus macaques and humans

**DOI:** 10.1101/2025.09.25.678525

**Authors:** Ugo Avila-Ponce de Leon, Shadisadat Esmaeili, Katherine Owens, Joshua T. Schiffer

## Abstract

Early during the COVID-19 pandemic, non-human primate (NHP) infection models emerged as highly useful tools for preclinical screening of antiviral drugs. However, it is uncertain whether NHP models can be used to precisely inform optimal dosing in humans. We previously established and validated mathematical models which were fit to SARS-CoV-2 viral loads from human clinical trials. These models identified that plasma drug concentrations required to inhibit viral replication by 50% in humans (*in* vivo EC50) differ substantially from *in vitro* EC50 estimates in cell culture systems. Here we apply models to sequential viral load data from SARS-CoV-2 infected rhesus macaques (RM) that were untreated or treated with nirmatrelvir/ritonavir, molnupiravir, or both drugs. We identify that equivalent plasma drug concentrations correspond to greater antiviral potency in lungs compared to nasal passages for nirmatrelvir and molnupiravir. Average nirmatrelvir antiviral efficacy in RM (30% in nasal passages and 46% in lungs) was estimated to be less than in humans (82%) due to shorter plasma drug half-life. Molnupiravir efficacy in RM (95% in nasal and 99% in lungs) is estimated to be similar to efficacy in humans against omicron variants. Our model estimates that 10-fold higher plasma nirmatelvir concentrations are needed in humans versus RM to achieve 50% reduction in viral replication, whereas 20-fold lower plasma molnupiravir concentrations are needed. Our results suggest that dose optimization in humans based on modeling of NHP viral loads is limited by drug-specific differences in pharmacokinetic, pharmacodynamic and virologic profiles, and that data from human phase 1 and 2 trials is better suited for this task.

## Introduction

Non-human animal models of infection are important for pre-clinical evaluation of antiviral agents.^1-12^ These models allow assessment of various metrics of drug efficacy including reduction in clinical severity, histopathologic tissue damage, and viral shedding.^13-18^ Because viral inoculum dose, site of infection, and timing are standardized across animals which are often from inbred colonies with similar genetic and immunologic histories, infected animals should in theory have less heterogeneous disease trajectories relative to humans. These factors may allow assessment of drug efficacy with a smaller sample size. An additional important advantage for SARS-CoV-2 is the ability to serially sample viral load and inflammatory damage from critical organs such as the lung which are rarely and inconsistently sampled in humans, and then only late during infection in a biased subset of critically ill patients.^19-21^

Non-human primates (NHP) are often employed to test antivirals due to similar anatomical and physiological properties between NHP and humans.^22-25^ SARS-CoV-2 infected NHP develop a self-limited respiratory illness like that of most humans with similar infection sites and types of immune responses.^1,7,12,26^ Early during the COVID-19 pandemic, most small molecular agents that were ultimately approved for human use were first tested in a variety of animal systems including ferrets,^27^ mice,^28-31^ and hamsters.^27,32-35^ Broadly speaking, NHP models sorted effective drugs such as molnupiravir,^36^ remdesivir,^24^ nirmatrelvir / ritonavir,^36^ and various monoclonal antibodies^37,38^ from ineffective drugs such as hydroxychloroquine.^39^

Yet there are important limitations to NHP models for testing antiviral drugs. The COVID-19 disease course in NHP is typically mild and duration of viral shedding is shorter than in humans with an earlier and lower peak viral load.^4,7,15,16,40^ Infection and treatment conditions also lack realism: rather than exposure by aerosolization,^18,41-43^ high titers of virus are usually directly inoculated onto nasal passage and airway mucosa.

The critical impact of stage of infection at the time of treatment initiation on treatment outcomes is also overlooked in most NHP studies. Treatment is nearly always implemented within 12 hours after infection during the pre-symptomatic phase which mimics post-exposure prophylaxis (PEP) rather than treatment of symptomatic disease.^24,44^ In humans, SARS-CoV-2 PEP has not been widely implemented relative to treatment.^45^ Drugs such as nirmatrelvir / ritonavir and remdesivir which are extremely effective as early symptomatic treatment failed as PEP, and were only partially effective as late symptomatic treatment.^45-54^ Additionally, prior to widespread vaccination, disease severity in humans was dictated by the intensity of immune responses 1-2 weeks after infection when viral levels were typically already in decline:^55-57^ NHP models fail to recapitulate aberrant immune responses which are characteristic of severe human infection. Another drawback of initiating treatment so early is the lack of concurrent immune responses which are vital to elimination of infected cells and work in synergy with antivirals, so the antiviral potential may be underestimated.^49,58^

Given the wealth of data from NHP models of infection, there is value in performing retrospective analyses to derive maximal value from these precious experiments. Mathematical models are a key tool for this purpose. To date, models have recapitulated outcomes of human SARS-CoV-2 clinical trials,^49,59-66^ and viral load and immune trajectories from NHP challenge experiments.^23,67,68^ The most accurate and realistic trial simulation models combine viral immune dynamic (VID), pharmacokinetic (PK) and pharmacodynamic (PD) equation sets and are calibrated to reproduce observed temporal reductions in viral load.^69,70^ Our group solved models for the human *in vivo* EC50, the plasma drug concentration required to inhibit viral replication by 50%, of nirmatrelvir/ritonavir and molnupiravir.^49,71^ This value is a critical target for plasma drug levels and for dose optimization in subsequent trials. We identified that for nirmatrelvir the *in vivo* EC50 is 13-100 fold higher than *in vitro* drug concentrations needed to inhibit viral replication by 50%.^49^ For molnupiravir, the *in vitro* EC50 is approximately 10 times higher than *in vivo* estimates.^71^ While these discrepancies are unsurprising given the extra physiologic processes which occur *in vivo* including turbulent blood flow, variable physiologic cell states, and dynamic plasma protein binding, a key question is whether mathematical models applied to viral load data from NHP models can arrive at more accurate estimates of the human *in vivo* EC50 compared to *in vitro* assays. Here, we estimate the *in vivo* EC50 of nirmatrelvir / ritonavir and molnupiravir in rhesus macaques (RM) using mathematical models and compare these results to estimates from humans.

## Results

### Antiviral effects of nirmatrelvir and molnupiravir in NHP nasal passage and lung

We used mathematical modeling to reproduce observed virologic outcomes in RM including data from 5 animals treated with nirmatrelvir/ritonavir (20 mk/kg + 6.5 mg/kg every 12 hours x 7 doses), 5 treated with molnupiravir (130 mg/kg every 1 hours x 7 doses) and 5 treated with both drugs (same doses).^7,36,72,73^ All treated RM were infected with doses of 2×10^6^ TCID50 of SARS-CoV-2 delta variant inoculated separately and split between the nasal passage and trachea. Treatment was initiated 12 hours after infection. Nasal swabs and bronchoalveolar lavage (BAL) samples were collected on days 1, 2 and 4 post-infection allowing quantification of virus in the nasal passages and lung respectively. Viral load was assessed with polymerase chain reaction (PCR) for total viral RNA and infectious viral load (V_TCID50_). All RM were immunologically naïve having never been previously infected or vaccinated.

To improve identifiability of the viral dynamic model, we included controls from the treatment study and other publicly available datasets. 43 controls were used for modeling infection off treatment and were from 4 studies including 30 animals infected with WA.1 strain, 5 infected with delta variant and 8 infected with omicron variant.^7,36,72,73^ Viral load was assessed serially after infection with PCR in all control animals but infectious V_TCID50_ was measured only for the 5 delta infected animals from the treatment study listed above.^36^

Overall, nirmatrelvir and combination treatment most potently reduced SARS-CoV-2 RNA in the nasal passages and lungs. Molnupiravir and combination treatment most potently reduced infectious SARS-CoV-2 levels. Relative to control (vehicle), the nasal passage of nirmatrelvirtreated animals had significantly reduced levels of viral RNA at days 2 and 4 **(Fig 1A)** and non-significantly reduced infectious virus at days 1 and 2 **(Fig 1B)**. In the lungs, nirmatrelvir-treated animals had significantly reduced levels of viral RNA at day 4 relative to control **(Fig 1C)** but did not have significantly decreased infectious virus **(Fig 1D)**.

**Figure 1.**
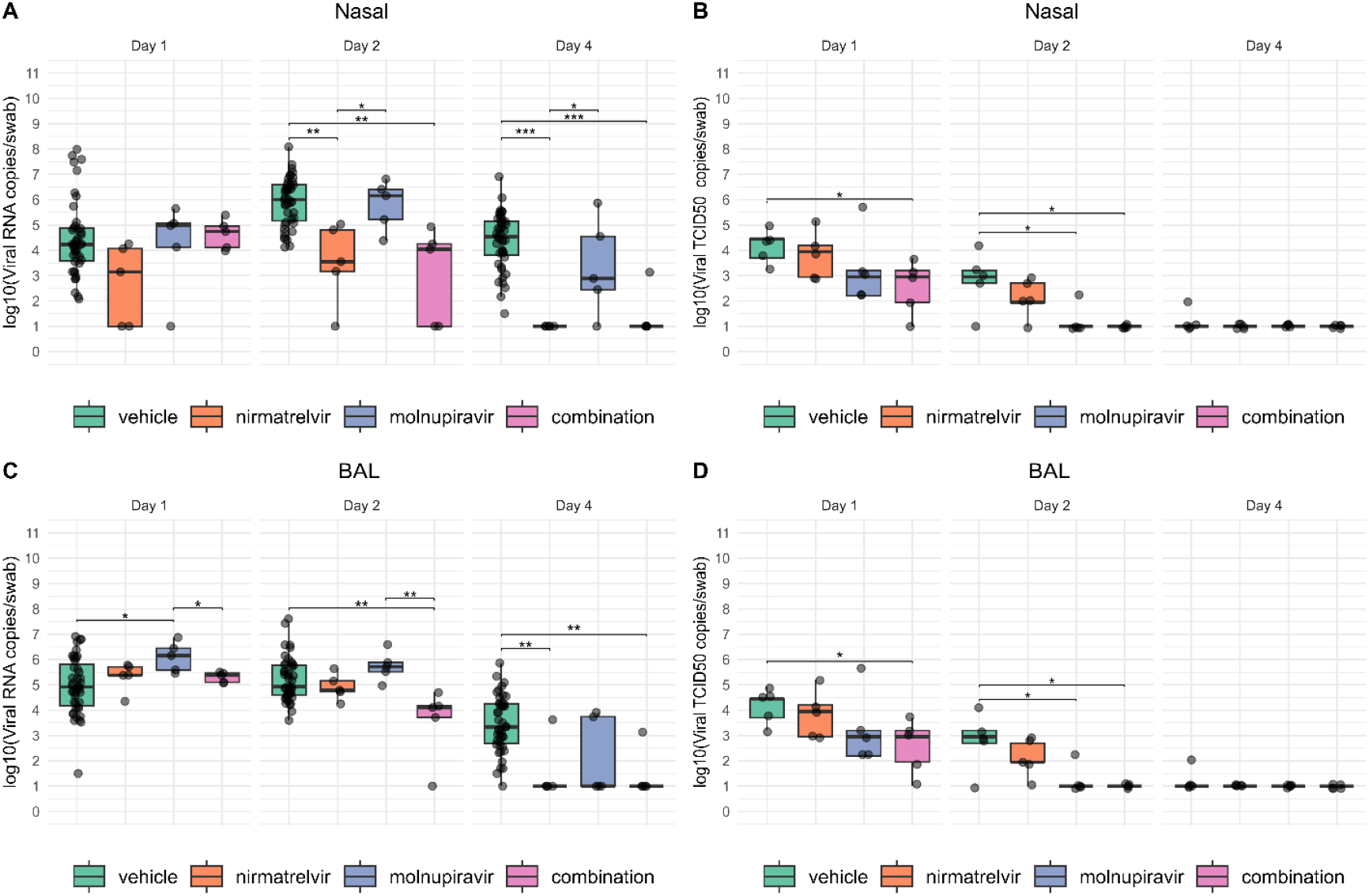
Observed viral loads during SARS-CoV-2 infection without treatment (n=43) treatment with nirmatrelvir (n=5), molnupiravir (n=5), and combination therapy (n=5). Nasal swabs and bronchoalveolar lavage (BAL) were collected at days 1, 2 and 4 post-infections in all animals and viral loads in each sample were quantitated using polymerase chain reaction and plaque assay. **(A)** SARS-CoV-2 RNA log10 copies/swab in the nose. **(B)** SARS-CoV-2 log10 V_TCID50_ / swab in the nose. **(C)** SARS-CoV-2 RNA log10 copies/mL BAL fluid in the lung. **(D)** SARS-CoV-2 V_TCID50_ / mL BAL fluid. Bonferroni corrected Wilcoxon test was carried out for the comparison between treatment arms. *p<0.05, **p<0.01, ***, p<0.001.

Relative to control (vehicle), the nasal passage of molnupiravir-treated animals had similar levels of viral RNA **(Fig 1A)** but infectious virus was non-significantly reduced at day 1, with significant reduction on day 2 **(Fig 1B)**. In the lungs, molnupiravir did not reduce levels of viral RNA compared to control **(Fig 1C)** but significantly reduced infectious virus at day 2, with a non-significant trend on day 1 **(Fig 1D)**.

Relative to control, combination nirmatrelvir and molnupiravir significantly reduced nasal levels of viral RNA at days 2 and 4 **(Fig 1A)** and infectious virus at days 1 and 2 **(Fig 1B)**. In the lungs, the combination reduced levels of viral RNA at days 2 and 4 **(Fig 1C)** and infectious virus at days 1 and 2 **(Fig 1D)**.

We observed high variability in viral RNA and infectious virus levels particularly at early timepoints, and in the control animals. Infectious virus was usually cleared from nasal passages: in controls, all animals except one cleared and the single outlier had a low TCID50 value, while all animals receiving nirmatrelvir and molnupiravir cleared infectious virus **(Fig 1B)**. A similar pattern was observed in lungs: in controls, all animals except one cleared and the single outlier had a low TCID50 value, while all animals receiving nirmatrelvir and molnupiravir alone or in combination cleared infectious virus **(Fig 1D)**.

Viral RNA was not eliminated from nasal passages or lungs in any controls by day 4 **(Fig 1A, 1C)**. In nasal passages, all animals treated with nirmatrelvir, and 1 out of 5 treated with molnupiravir cleared viral RNA **(Fig 1A)**. Viral RNA was eliminated from the lungs in 4 out of 5 animals treated with nirmatrelvir, and 3 out of 5 treated with molnupiravir **(Fig 1C)**. The heterogeneity of viral trajectories suggests that modeling of individual trajectories might be useful to explain inter-animal differences as well as treatment effects.

WA.1 and delta virus viral RNA kinetics appeared similar except for higher viral RNA for delta variant at days 1 and 2 in lung **(Fig S1)**. We therefore included the delta variants and WA.1 in the control arm to increase sample size for modeling.

### Mathematical model for SARS-CoV-2 viral and immune dynamics of untreated infection in non-human primates (NHP)

We first developed a mathematical model to accurately simulate viral load dynamics and estimate viral and immune parameters in the absence of therapy. We based the general framework on our model of human SARS-CoV-2 infection which was validated against viral load data from 1510 untreated infections.^58^ We elected to use this model as our baseline to allow parallel estimation of drug potency in humans and RM. However, for our NHP model we did not include an acquired immune response, as virus was often cleared prior to this response becoming relevant.

Given the added presence of TCID50 data which was not available for our human modeling, we extended the model from Owens et al. to estimate infectious virus levels over time. We evaluated a total of 8 computational models, each incorporating different mechanistic assumptions considered critical to accurately capture viral dynamics. Specifically, we tested models assuming that infectious virions are cleared either at the same rate as productively infected cells or at the same rate as the total viral population. In addition, we examined the impact of including or excluding refractory and eclipse cell compartments. Refractory cells, which are temporarily resistant to infection due to innate immune responses or drug effects, can significantly alter viral spread and treatment outcomes. Similarly, the eclipse phase during which cells are infected but not yet producing virus can introduce important delays in viral production, affecting model predictions of viral load kinetics and rebound. These comparative analyses aimed to determine which biological features most critically shape observed infection and treatment response patterns.

Among the models tested the optimal fit to experimental data based on the lowest Akaike Information Criteria (AIC) score **(Fig 2A)** included both refractory and eclipse compartments and assumed that infectious viruses were cleared at the same rate as infected cells **(Table S1)**. The model is therefore highly aligned with our human trial simulation model.

**Figure 2.**
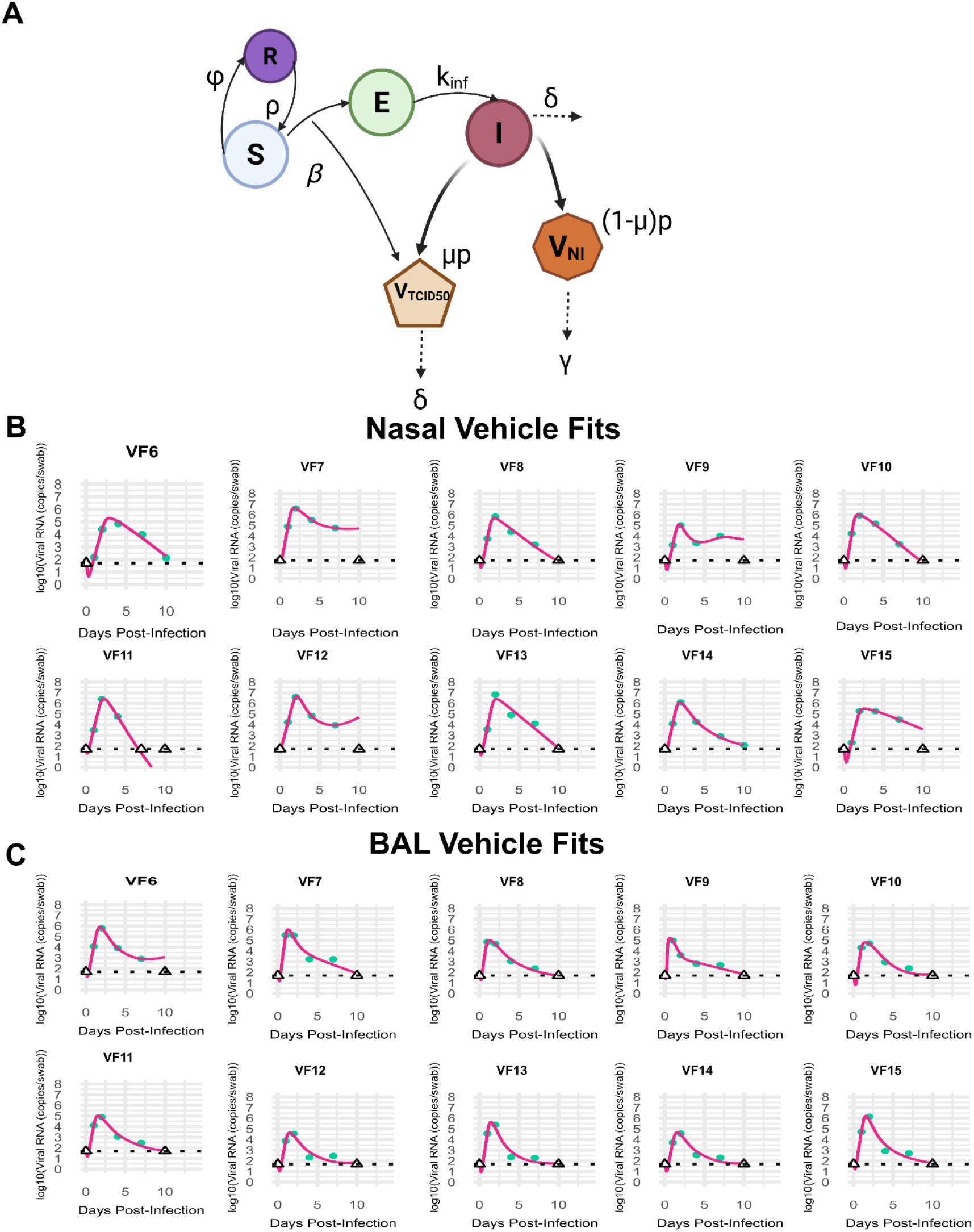
Optimized SARS-CoV-2 Viral Immune Dynamic Model. **(A)** Schematic of the system of ordinary differential equations to model the viral dynamics with state variables in capital letters: S: susceptible targets cells, E: Eclipse infected cells, I: infected cells, R: Refractory cells, V_TCID50_: Infectious viral particles, V_NI_: non-infectious virus. V_RNA_ is captured by PCR and is the sum of V_TCID50_ and V_NI_. **(B)** Model fits (pink lines) to observed SARS-CoV-2 V_RNA_ (pink dots) in the nasal compartment. **(C)** Model fits to observed SARS-CoV-2 V_RNA_ in the BAL compartment. The dashed black line in panel B is the limit of detection. Hollow triangles overlaying the line are undetectable viral load.

Our optimal model assumes a finite number of susceptible cells (S) that get infected via a force of infection, β, by infectious viral particles (V_TCID50_). When a cell is infected, it enters an eclipse phase © that lasts an average of 1/k_inf_ days before subsequent entry into a productively infected phase (I). Productively infected cells produce virions at rate *p*. Most of viral RNA is non-infectious (V_NI_), but a small proportion, *μ*, is infectious and constitutes TCID50 (V_TCID50_). For the purpose of comparison with data, V_RNA_ = V_NI_ + V_TCID50_. Given that most infection occurs via cell-to-cell spread of infectious viral particles, we assumed V_TCID50_ remains inside the cell and decays according to the lifespan of the infected cell with rate δ. However, since viral RNA persists outside of cells, in our model V_RNA_ decays at a separate clearance rate, *c*.

In the model, susceptible cells that encounter productively infected cells temporarily become refractory to infection with the rate *ϕ* and revert back to the susceptible state at rate *ρ*. This agrees with our human model and other models validated against human data^74^. This mechanism can be interpreted as innate immunity because its effect correlates directly with intensity of infection and assumes no immunological memory (**Fig 2A**). We employed the same model structure for nasal and BAL sites but allowed different parameter estimates.

### Mathematical model fit to untreated and infected RM

Using a mixed-effects population fitting approach implemented in Monolix, we estimated parameter values that achieve good fits to individual viral load data from 43 NHP quantified with nasal PCR (**Fig 2B, Fig S2**) and bronchoalveolar lavage (BAL) PCR (**Fig 2C, Fig S3**). The model simultaneously captured levels of infectious and total virus from the 5 NHP with TCID50 quantitation in nasal passage and BAL (**Fig S2, Fig S3**). For 9 RM of the nasal passage (VF7, VF9, VF12, VF18, VF20, VF40-VF43) and 8 RM for the lung compartment (VF6, VF17, VF23, VF29, and VF40-VF43), the model does not predict viral elimination, though all animals except VF40 appeared to be in the process of viral clearance at the time of the last sample. The treatment period only extends from day 0.5-4 post infection, a timeframe in which the model matches the data closely. Projections later during infection, when the acquired immune response may be active, and beyond the observation period are less reliable.

We next compared estimated parameter values between the two anatomic compartments (**Table 1**). Infected cell death rates were slightly lower in the lung, while viral replication rate was slightly higher in the nasal passages compared to the lungs in the untreated animals. The viral infection rate was higher in the lung compared with the nasal passages, and a higher proportion of viral RNA was predicted to be infectious in the lung. The conversion rate of cells from susceptible to refractory was slightly higher in lung while waning of this protection was also higher in lung.

**Table 1.** Population parameters obtained from our mathematical model for nasal and lung compartments.

| Parameters (units) | Nasal (standard error) | Lung (standard error) |
| --- | --- | --- |
| $\beta \log_{10}(PFU/ml)^{-1}day^{-1}$ | $10^{-3.22}$ (0.19) | $10^{-2.42}$ (0.15) |
| $\delta$ ($days^{-1}$) | 4.62 (0.65) | 4.16 (0.51) |
| $\mu$ | 0.0084 (0.0042) | 0.02 (0.012) |
| $p$ ($\log_{10} days^{-1}$) | $10^{1.76}$ (0.17) | $10^{1.17}$ (0.16) |
| $\phi$ ($\log_{10} days^{-1}$) | $10^{-4.74}$ (0.37) | $10^{-4.29}$ (0.22) |
| $\rho$ ($\log_{10} days^{-1}$) | $10^{-2.47}$ (0.66) | $10^{-2.06}$ (0.37) |

We next compared parameter ranges between viral variants (WA.1 (in color blue), delta (in color red), omicron (in color green)) for each anatomic compartment and observed that several key virological parameters differ **(Table S2)**. The infectivity rate (β) was significantly higher for omicron than WA.1 and delta across compartments, aligning with its known increased transmissibility. Infected cell death (δ) was significantly larger for delta in both sites, particularly in BAL, suggesting greater cytopathology. Infectious cell proportion (*μ*) was highly variable, but omicron often showed higher median values, possibly reflecting enhanced cell entry. Viral production (p) was notably higher for delta in the nasal compartment, whereas omicron peaked in BAL. Innate immune evasion (*ρ*) was larger (weaker protection) for omicron, especially in BAL, consistent with its immune escape traits. The rate of conversion to a protected state (φ) was slower for WA.1, while Omicron infection resulted in faster refractory conversion particularly in BAL, suggesting shorter-lived innate immune responses. These differences underscore variant-specific adaptations in transmission, virulence, and immune interaction across respiratory niches.

### Modeling nirmatrelvir treatment in NHP

We next extended our model to simulate viral load dynamics in RM treated with nirmatrelvir. We modeled treatment with the assumption that nirmatrelvir inhibits viral replication (*p*) with efficacy (*ε*). Efficacy over time is determined by combining PK and PD models.

We used a two-compartment pharmacokinetic (PK) model (**Fig 3A**) to accurately reproduce plasma drug levels in 18 uninfected NHP that received nirmatrelvir dosed at 20 mg/kg, 50 mg/kg or 300 mg/kg twice daily, 6 and 18 apart, for fifteen days.^75^ As depicted in **Figs S4-S6**, the model accurately reproduced plasma drug levels at the start and completion of the dosing cycle. Parameters are in **Table S3**. While peak drug level following 20 mg/kg dosing in NHP were close to that in humans (∼3000 ng/mL), we noted a considerably lower nirmatrelvir elimination half-life in RM from our data (∼ 3 hours) relative to ∼6 hours in humans.^49^ As a result, drug troughs in humans with twice daily dosing are ∼500 ng/mL whereas in RM, they are projected to be <10 ng/mL.

**Figure 3.**
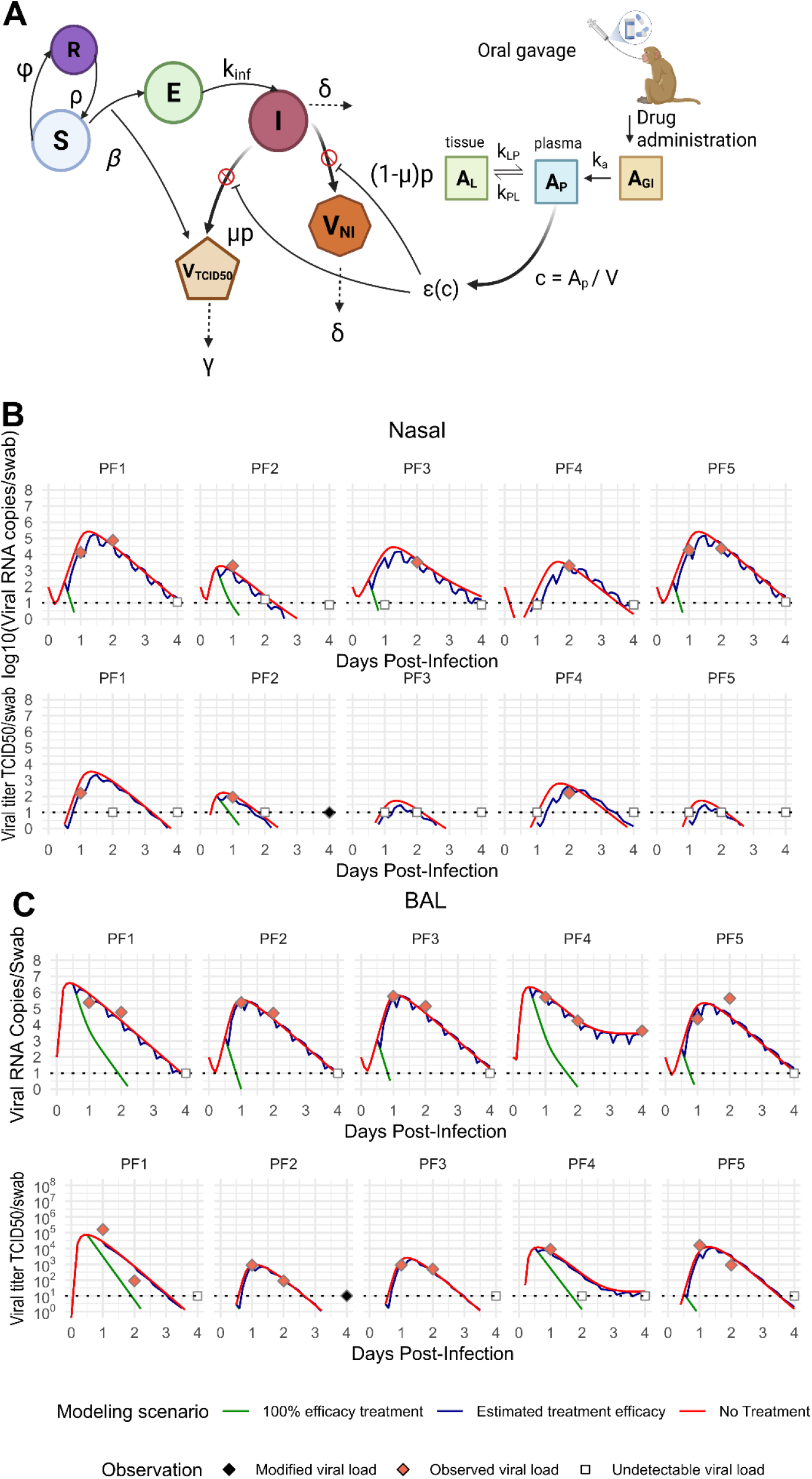
Mechanistic mathematical model with fits to viral load in nirmatrelvir treated RM. **(A)** Schematic of the ordinary differential equations to model viral dynamics. PK model and drug activity are also shown. **(B)** Model fits to nasal passage SARS-CoV-2 RNA (row 1) and V_TCID50_ data (row 2). **(C)** Model fits to BAL SARS-CoV-2 RNA (row 1) and V_TCID50_ data (row 2). In all panels of B and C, the dashed black line in is the limit of detection. Hollow squares overlaying the line are undetectable viral load. Orange diamonds are observed viral loads. Black diamonds are modified viral loads (see Methods). Blue lines are estimated model projections given true efficacy. Red line is counterfactual projection with no drug. Green line assumes 3-day treatment with a fully effective drug.

To measure efficiency of the drug based on the concentration we used a hill equation to model drug pharmacodynamics (PD).^76 77,78^ We used the parameters obtained from in vitro efficacy data collected at different doses of nirmatrelvir. The *E*_*max*_, the invitro *EC*_50_ and the hill value *n* were fixed based on the values estimated in the experiment performed to evaluate the efficacy of nirmatrelvir in Calu-3 cells infected with the delta variant. The in-vivo *EC*_50_ was estimated based on the potency adjustment factor defined as 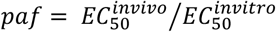. The *E*_*max*_, Hill coefficient (n), and *EC*_50_ values derived from human Calu-3 cells are considered pharmacodynamically relevant for non-human primate (NHP) studies, as these parameters reflect the intrinsic activity of the compound at a conserved biological target. Calu-3 cells, which model human airway epithelium, express key host factors and cellular pathways that are functionally similar in NHPs. Given the high degree of conservation in drug target engagement and downstream antiviral mechanisms across species, in vitro PD parameters from human cell-based assays can serve as reliable surrogates for in vivo PD modeling in NHPs. This translational approach is particularly justified when the mechanism of action is host-directed and less susceptible to species-specific differences.

We fit the integrated viral dynamic and PK-PD models to the SARS-CoV-2 PCR and TCID50 data from five rhesus macaques infected with 2 × 10^6^ TCID50 (delta variant) that received nirmatrelvir dosed 20 mg/kg for 4 days^36^. Viral loads were assessed at day 0, 1, 2 and 4 of treatment in upper airway (**Fig 3B**) and the lung (**Fig 3C**).^36^ Population PK and PD parameters were used to simulate nirmatelvir treatment, but individual viral dynamic parameters were estimated along with an in vivo EC50 shared acrodd animals. Model simulations reproduced the observed PCR and V_TCID50_ data (blue lines, **Fig 3B & 3C**).

By performing counterfactual simulations assuming the five treated animals did not receive treatment (*ε* = 0) and by assuming treatment with perfect efficacy (*ε* = 1), we identified that nirmatrelvir had an observable but incomplete effect on viral load reduction (red and green lines respectively, **Fig 3B & 3C**). Treatment effects on V_TCID50_ typically paralleled those of V_RNA_.

Viral load reduction patterns during treatment more closely approximated no efficacy than full efficacy. Nevertheless, viral reductions relative to no treatment at peak drug levels were often 0.5-1.0 log. Moreover, reductions in viral area under the curve (a surrogate for infected cell surface area in viral dynamic models)^79^ during the treatment window were significant. In nasal samples, treatment resulted in a median viral PCR (between day 0.5 and 3.5) AUC reduction of 43.4% (range: −10.2% to 56.8%) and a 43% reduction in V_TCID50_ (range: −33.5% to 56.8%) relative to control (**Fig S7A & B**). In BAL (between day 0.5 and 3.5), viral decline was slightly more significant and less variable. We a project a median viral PCR AUC reduction of 55.4% (range: 49.4–57.1%), and a 58.5% reduction in V_TCID50_ (range: 56.7–59.5%) (**Fig S7C & D**).

### Nirmatrelvir antiviral potency in nasal and lung cells

We combined PK and PD models with the solved value for *in vivo* EC50 to estimate the instantaneous values of drug efficacy over time. Due to parameter identifiability issues, we were forced to estimate a single plasma *in vivo* EC50 value for all animals though separate values were estimated for nasal passage and lung.

Assuming equivalent PK for each animal, we estimated drug efficacy as a function of time **(Fig 4A, B)**. Efficacy was nearly complete at the time of drug peak but rapidly decreased between doses due to short nirmatrelvir half-life. The time-averaged efficacy during treatment was 30% and 46% for the nasal and lung compartments respectively. The slightly better drug activity in lung versus nasal tissues for the drug in NHP was due to slower initial decrease from peak drug level and higher antiviral activity at drug trough **(Fig 4A, B)**. Notably, our prior estimate for nirmatelvir / ritonavir efficacy in humans was 82% in nasal and oral compartments.

**Figure 4.**
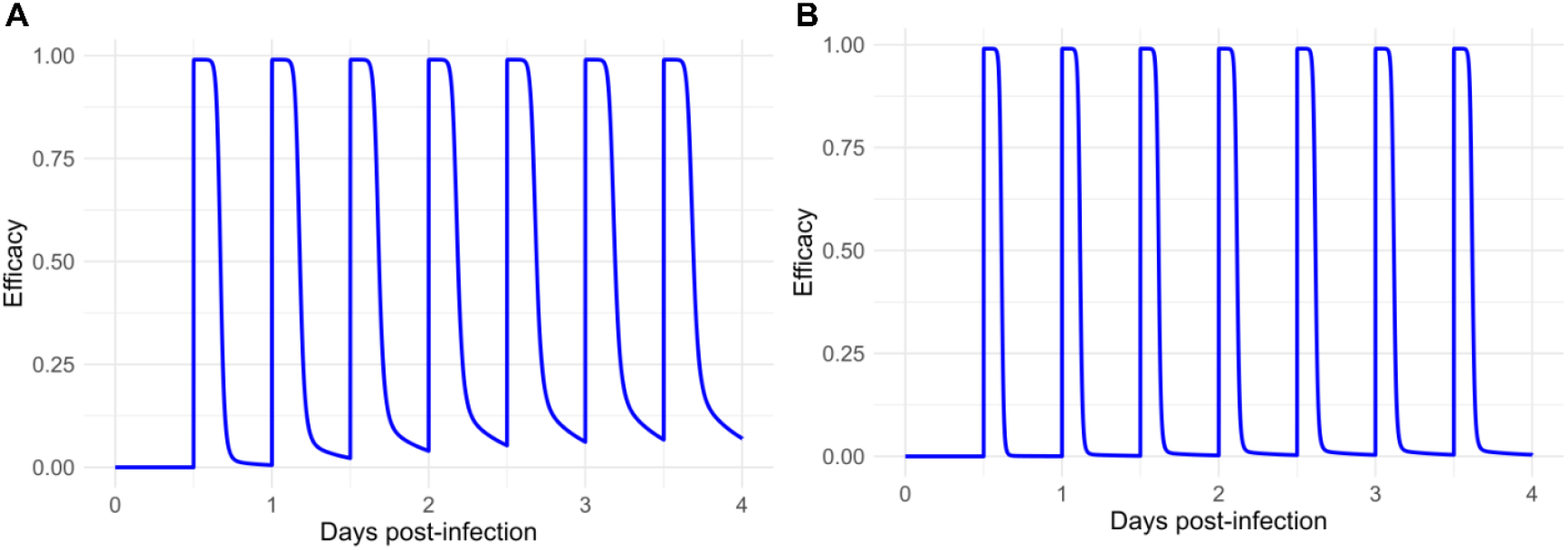
Model estimated efficacy of nirmatrelvir in different anatomic compartments. (A) Dynamic drug efficacy in the nasal compartment. (B) Dynamic drug efficacy in the lung.

### Modeling molnupiravir treatment in NHP

We also used our model to simulate viral load dynamics in RM treated with molnupiravir. Based on molnupiravir’s mechanism of action of viral error induction which terminally mutates the virus,^80^ we mimicked our human model by adding a new compartment to our model labeled mutated virus or *V*_*mut*_.^71^ To estimate the treatment efficiency of molnupiravir, we assumed the drug induces infected cells to produce *V*_*mut*_ at drug efficacy *ε*. The remaining produced virus (1-*ε*) is then divided into V_NI_ (1-*μ*) or V_TCID50_ (*μ*). Like V_NI_, *V*_*mut*_ is not infectious but is still detected with polymerase chain reaction (PCR) because the chance of a drug-induced mutation occurring in the PCR primer is less than 1% **(Fig 5A)**. For the molnupiravir model V_RNA_ = V_TCID50_ + V_NI_ + *V*_*mut*_.

**Figure 5.**
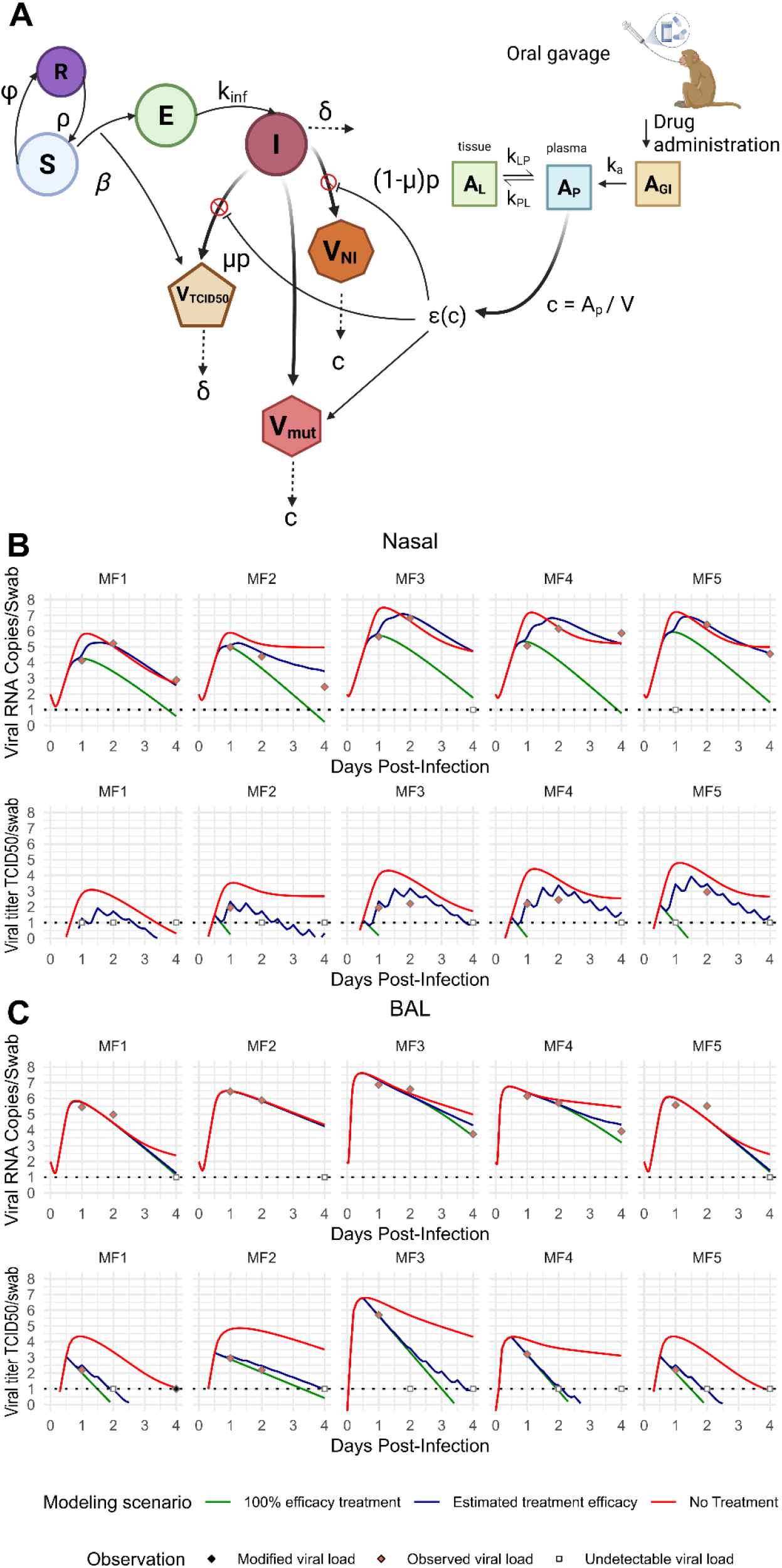
Mechanistic mathematical model with fits to viral load for molnupiravir treated RM. **(A)** Schematics of the system of ordinary differential equations to model the viral dynamics. The molnupiravir mechanism of action is to induce multiple errors during replication which means that when the concentration is sufficiently high, infected cells produce more mutated non-infectious virus rather than typical virions. Drug levels and activity are also shown. **(B)** Model fits to SARS-CoV-2 PCR and V_TCID50_ data from nasal compartment **(C)** Model fits to SARS-CoV-2 PCR and V_TCID50_ data from BAL compartment. The dashed black line in panel B is the limit of detection. Hollow squares overlaying the line are undetectable viral load. Orange diamonds are observed viral loads. Blue lines are estimated model projections given true efficacy. Red line is counterfactual projection with no drug. Green line assumes 3-day treatment with a fully effective drug.

We captured the PK of molnupiravir with a two-compartment model **(Fig S9)** to accurately reproduce plasma drug levels in 8 uninfected NHP that received molnupiravir dosed at a single dose of 100 mg/kg and was measured at different hours during a twenty-four hour time point.^75^ Peak drug levels (∼3000 ng/mL) following 100 mg/kg dosing approximated levels in humans following dosing with 800 mg. Drug trough was usually <10 ng/ml suggesting a rapid half-life of ∼3 hours. PK parameters are in **Table S3**. To measure efficiency of the drug based on the concentration we used a hill equation approach to model drug pharmacodynamics (PD) and we fixed the parameters for *E*_*max*_, the invitro *EC*_50_ and the hill value *n* that were parametrized based on experiments that measured the efficacy of molnupiravir in Calu-3 cells infected with the delta variant because they resemble the same pathway of infection as in the case for nirmatrelvir. The only value we estimated was the in-vivo *EC*_50_ based on the value of the potency reduction factor.

We fit the combined viral dynamic PK-PD model to nasal **(Fig 5B)** and lung **(Fig 5C)** data from 5 molnupiravir-treated animals, simultaneously matching both PCR (V_TCID50_ + V_NI_ + V_mut_) and V_TCID50_. As with nirmatrelvir, we used population PK and PD parameters to simulate molnupiravir treatment but estimated individual viral dynamic parameters along with a shared in vivo EC50. In keeping with the data, our model projects high antiviral potency in the nasal compartment (demonstrated by variably reduced peak V_RNA_ and substantial decrease in V_TCID50_). In lung, V_TCID50_ projections approximated a fully efficacious drug. In both cases, V_TCID50_ levels more rapidly decreased relative to V_RNA_. This is because PCR detected persistent high levels of mutated viral RNA in nasal **(Fig 6A)** and lung **(Fig 6B)** compartments. In the nasal passages and lungs of all animals, the majority of detectable SARS-CoV-2 viral RNA was drug mutated at all time points after the initiation of molnupiravir.

**Figure 6.**
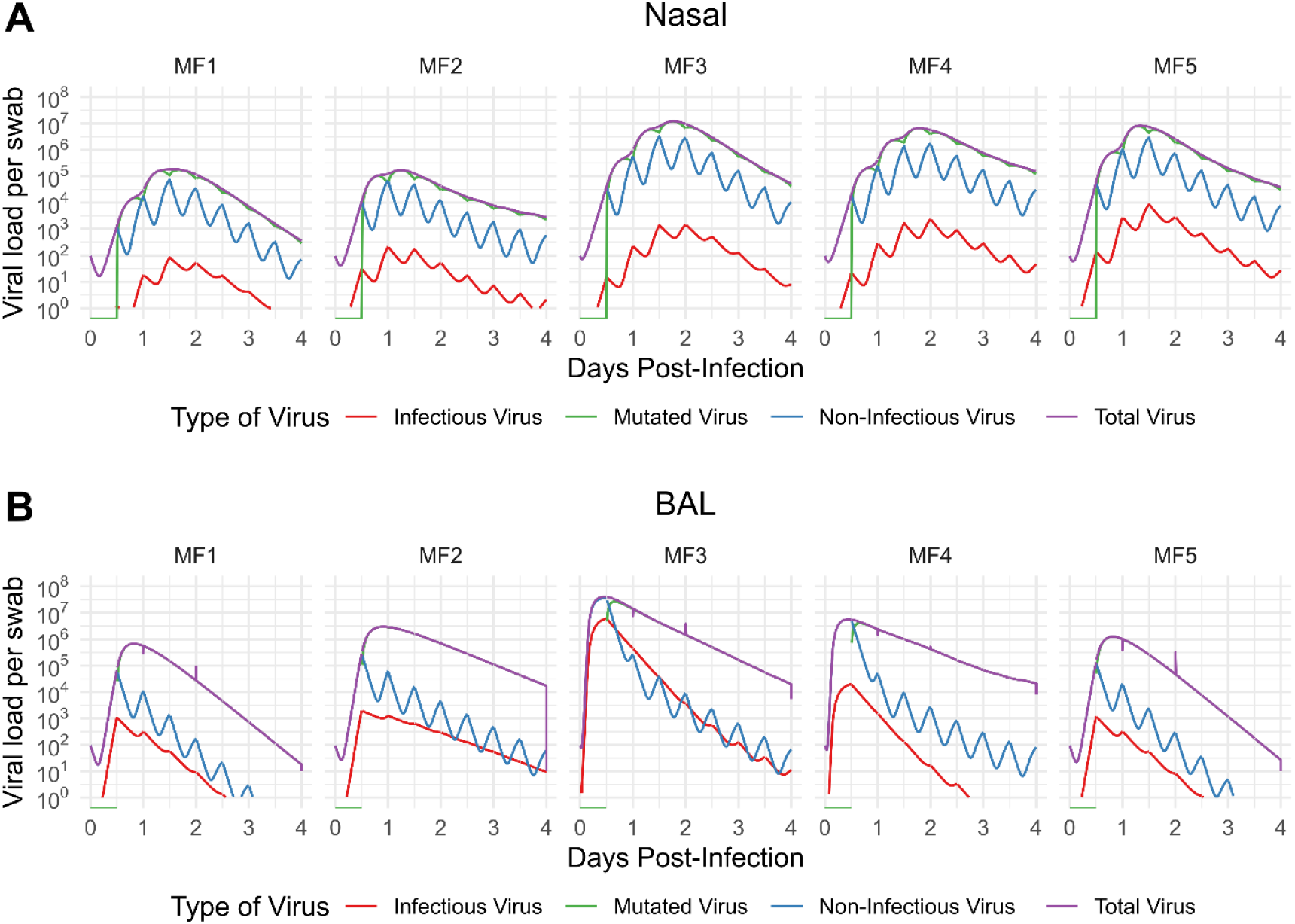
Dynamics of drug mutated viral RNA during molnupiravir treatment. For the 5 molnupiravir-treated animals, we plotlet the simulated viral load stratified by the model-predicted category of virus: V_TCID50_ (red), V_mut_ (green), V_NI_ (blue) and V_RNA_ (purple) in both the (A) nasal passage and (B) the lungs. All lines are an output from our mathematical model, shown compared with data in **Fig 5**. In both compartments, most V_RNA_ is V_mut_ due to high drug potency.

We again performed counterfactual simulations assuming the five treated animals did not receive treatment (*ε* = 0) and by assuming treatment with perfect efficacy (*ε* = 1). Notably, molnupiravir resulted in significant reductions in viral area under the curve (a surrogate for infection surface area) for all animals in all compartments compared to counterfactual simulations of no treatment. Molnupiravir exhibited distinct antiviral effects across compartments: In nasal samples, it achieved a median 46.7% reduction in viral RNA (between day 0.5 and 3.5) (PCR; range: 36.1–62.1%) and a 94.2% reduction in infectious virus (V_TCID50_; range: 93.3–95.6%), demonstrating potent activity in the upper respiratory tract. In contrast, bronchoalveolar lavage (BAL) revealed compartment-specific variability—while viral RNA (PCR) showed only modest declines (median: 5.2%; range: 4.7–13.4%), infectious virus (V_TCID50_) was robustly suppressed (median: 98.3%; range: 81.6–98.4%). This dissociation suggests molnupiravir’s mechanism (lethal mutagenesis) preferentially targets replicating virus over pre-existing RNA in the lower airways).

### Molnupiravir antiviral potency in nasal and lung cells

We combined the PK and PD models to estimate instantaneous values of molnupiravir efficacy over time. Due to parameter identifiability issues, we estimated a single value for plasma *in vivo* EC50 across all animals though separate values were estimated for nasal and BAL. Assuming equivalent PK for each animal, we estimated drug efficacy as a function of time **(Fig 7A, B)**. In nasal passages, efficacy was nearly complete at the time of drug peak but decreased somewhat at drug trough due to short half-life. In lung, the estimated efficacy was nearly complete throughout the dosing interval. The time-averaged efficacy during treatment was 95% and 99% for the nasal compartment and lung respectively, indicating better drug activity in lung versus nasal tissues for the drug in NHP.

**Figure 7.**
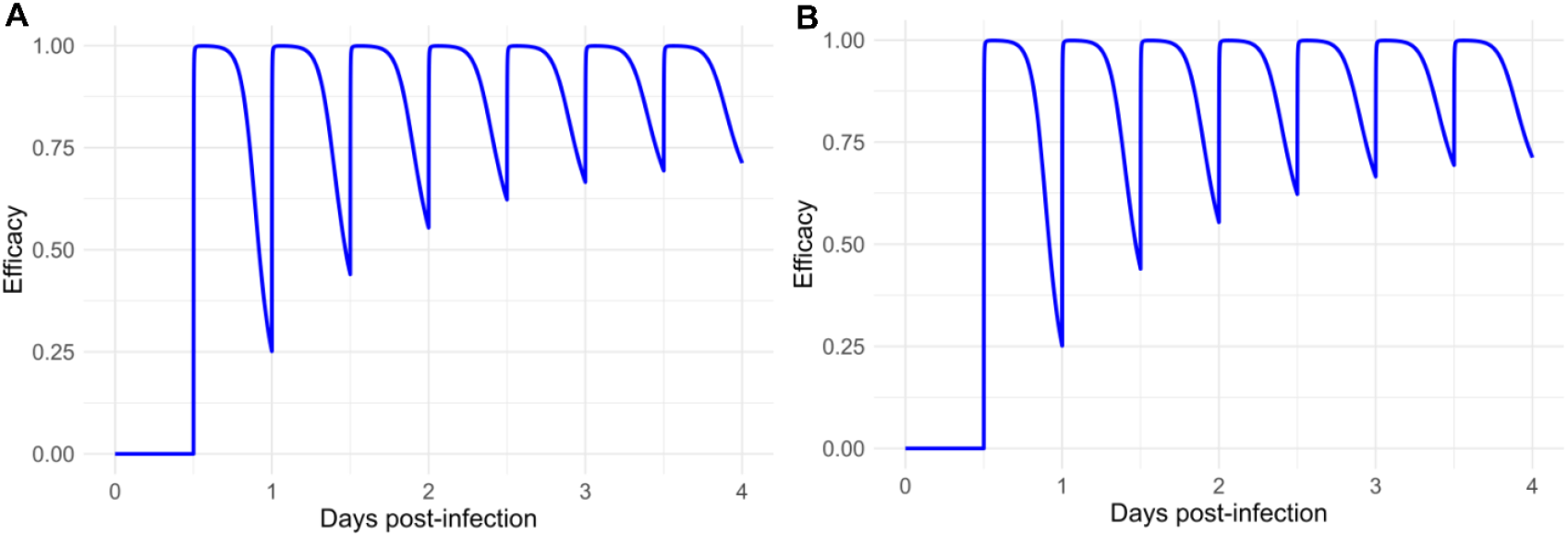
Estimated efficacy of molnupiravir in different anatomic compartments. (A) Dynamic drug efficacy in the nasal compartment. (B) Dynamic drug efficacy in the lung.

### Comparison of estimated in vivo EC50 across species

We next compared estimated in vivo EC50 for NHP with human and in vitro values. For molnupiravir, the estimated plasma concentration of drug required to limit replication by 50%, in nasal passage and BAL, was less than *in vitro* estimates but still higher than required plasma concentrations in humans; molnupiravir was predicted to be 7 times more potent in NHP lung than in nasal passages **(Fig 8A and Table 2)**. For nirmatrelvir, the estimated plasma concentration of drug required to limit replication by 50% was similar to current *in vitro* estimates but ∼10-fold less than human plasma concentrations; nirmatrelvir was predicted to be 2 times more potent in NHP lung than in nasal passages **(Fig 8B and Table 2)**. Notably, all of our human estimates were derived from trials conducted during the omicron variant wave, whereas NHP estimates were with the delta variant and *in vitro* estimates were with multiple variants. Overall, these results suggest that *in vivo EC*_50_ values estimated from fitting models to NHP viral load data are insufficient for projecting drug potency in humans.

**Figure 8.**
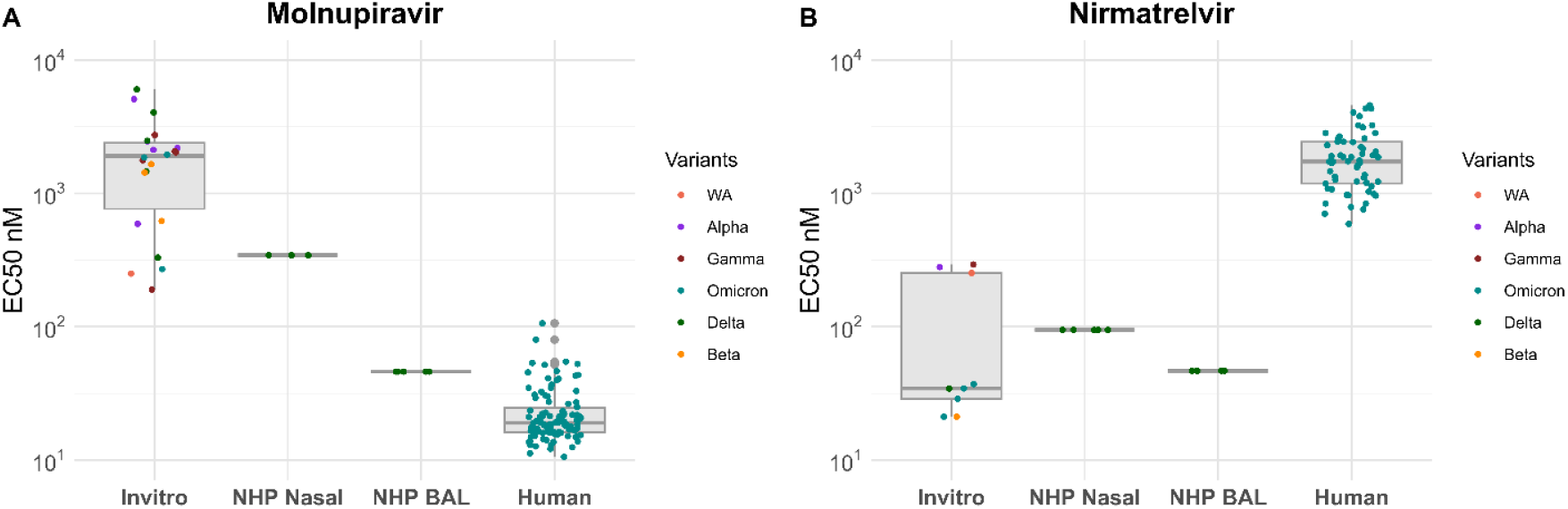
Estimated EC50 in NHP versus humans. Comparison of the values of EC_50_ for in vitro studies and the values estimated from our mathematical model in NHP and humans for **(A)** molnupiravir and **(B)** nirmatrelvir.

**Table 2.** Summary of antiviral activity for Molnupiravir and Paxlovid across different systems. log10 EC50 from figure 8 are shown. Median, interquartile range (IQR), and 5–95% ranges are shown for in vitro experiments, non-human primate (NHP) nasal and BAL samples, and human data.

| Molnupiravir |  |  |  |  |
| --- | --- | --- | --- | --- |
|  | in vitro | NHP Nasal | NHP BAL | Human |
| Median | 3.27 | 2.53 | 1.66 | 1.27 |
| IQR | 3.19 | 0 | 0 | 0.91 |
| 5-95% | 2.39-3.7 | 2.53-2.53 | 1.66-1.66 | 1.1-1.7 |
| Paxlovid |  |  |  |  |
| Median | 1.53 | 1.97 | 1.66 | 3.23 |
| IQR | 1.35 | 0 |  | 3.09 |
| 5-95% | 1.32-2.45 | 1.97-1.97 | 1.66-1.66 | 2.89-3.61 |

### Combination treatment in NHP

As a model validation procedure, we used the model identified *in vivo* EC50 value for both drugs and fit the model to the observed nasal and lung viral loads in RM treated with both drugs **(Fig S10)**. As with monotherapy, we used the population PK-PD parameters for both drugs and estimated individual viral dynamic parameters for each animal. The model reproduced nasal V_RNA_ and V_TCID50_ in nasal passages **(Fig S10A & B)** and lungs **(Fig S10C & D)**. Both drugs alone and in combination had similar projected results on nasal V_RNA_, but molnupiravir was responsible for more of the observed decline of V_TCID50_ than nirmatrelvir. Most V_RNA_ decline in lung resulted from nirmatrelvir as molnupiravir converted V_NI_ to V_mut_ which is detectable with PCR. Overall, the individual effects of each drug were evident in the observed viral kinetics on combination treatment.

## Discussion

During the COVID-19 pandemic, NHP models of infection accelerated antiviral development by proving that certain drugs dramatically limit clinical severity, lower viral load, and decrease histopathologic damage in lung.^24,36-38,68,81^ Given the expense and difficulty of these experiments, it is important to maximally extract relevant conclusions from this data. We applied mathematical models to viral load data to assess whether *in vivo* drug potency in NHP approximates that observed in humans, and whether this information could be leveraged for dose optimization in humans. Our models benefitted from the additional data from NHP including concurrent measurement of SARS-CoV-2 RNA by PCR and TCID50 by culture in upper airways and lungs but were challenged by small sample size and noise in the data.

Our optimal models reproduced both viral RNA and TCID50 in nasal passages and lungs under treatment. We obtained estimates of the *in vivo* EC50 which suggest that the plasma concentration required for molnupiravir and nirmatrelvir differs between NHP and humans. Whereas higher molnuporavir plasma concentrations are required to limit SARS-CoV-2 replication in NHP relative to humans, the opposite appears to be true for nirmatrelvir. While our NHP estimates of *in vivo* EC50 in lungs for molnupiravir better approximate those obtained by our modeling of human data, they are insufficiently accurate for projecting dose selection in human trials. Overall, these results suggest that data from phase 1 and 2 human clinical trials should be leveraged for dose selection in phase 3 trials, which in turn implies that phase 1 and 2 trials should be designed with sufficiently granular PK and virologic sampling to facilitate accurate modeling.

Surprisingly, nirmatrelvir had lower time averaged antiviral efficacy in RM relative to humans despite the need for less drug in plasma to inhibit viral replication by 50%. This is due to the extremely rapid clearance of nirmatrelvir in RM relative to humans resulting in a significant portion of the dosing interval with sub-therapeutic levels. Molnupiravir PK and PD properties are slightly more similar between humans and RM. We previously estimated molunpiravir efficacy of 53% against the delta variant and 95% against omicron in humans,^71^ whereas in RM time-averaged efficacy was 95% and 99% effective against delta in nasal passages and lungs respectively. Differences in efficacy between RM and humans may relate to differences in weight-adjusted dose, PK properties, drug protein binding, and cellular concentration needed to inhibit viral replication.

We also identified that higher plasma molnupiravir concentrations are required for both drugs in NHP nasal passages relative to lungs. It is unknown whether a similar relationship exists in humans as lung viral loads are almost never measured. This result suggests the possibility that drug concentrations which do not eliminate upper airway shedding are sufficient to limit extent of infection in the lung.^23^ Viral load reduction in upper airways was established as a valid surrogate endpoint for reduction in hospitalization by comparing results across trials.^82^ Our results suggest that a drug that does not reduce upper airway viral loads could in theory be associated with benefit if there are more potent effects in the lung. Sampling of lung with BAL is a significant advantage of animal models over human infection trials when assessing drug potency.

Our modeling of nirmatrelvir is relatively straightforward based on a simple mechanism of action. Molnupiravir converts viral RNA to a mutated form which is still largely detectable with PCR.^71^ This requires a different model than protease inhibitors like nirmatrelvir and explains why molnupiravir disproportionately lowers TCID50 but not PCR detectable viral RNA in both RM and humans. This result reinforces our previous prediction that standard viral PCR underestimates the true potency of molnupiravir.^71^ Our model suggests that molnupiravir is far more potent than nirmatrelvir in RM. If a more favorable half-life was assumed for nirmatrelvir, then both drugs would be highly potent.

This modeling exercise has significant limitations mostly related to the nature of available data. Despite advantages of controlling infection conditions, standardization of infection dose, measuring both viral PCR and culture, and sampling both lung and nasal passages, the data from these animals is at the limit of what is possible to model. The frequency and duration of sampling and the number of infected animals was low relative to human trials. We encountered parameter identifiability issues and had to estimate a single *in vivo* EC5O value across animals, which may not be realistic. We assumed equivalent drug PK for each animal by imputing population parameter values, which is an oversimplification. We also detected noise in the data and findings that were hard to rationalize biologically, including one timepoint with detectable TCID50 but negative PCR. For this reason, we had to make several subjective decisions about data censoring which would not have been necessary in a larger, more granularly sampled cohort. We performed counterfactual simulations to quantify the estimated effects of perfectly effective drugs, but these were not validated against larger existing datasets as we have done in humans, again owing to limited sample size. Along with estimated EC50 values that differ from human estimates, these key limitations further support our conclusion that modeling of viral load data from NHP is not sufficient for precisely informing human antiviral dosing strategies.

In summary, we fit mathematical models to a diverse dataset of SARS-CoV-2 infection including serial measures of viral load measured with PCR and TCID50 in both nasal passages and lung. We identified moderate drug efficacy for nirmatrelvir in both compartments due to short half-life in RM, and potent activity of molnupiravir, particularly in lung. This potent effect was only observed by modeling TCID50 rather than PCR as drug mutated virus persists on therapy. Despite the high utility of NHP models for broadly identifying agents with high therapeutic potential, our models suggest that estimates of plasma efficacy in NHP do not sufficiently predict efficacy in humans to inform dose selection.

## Material and Methods

### Preclinical data

We analyzed viral load observations from nasal and bronchoalveolar (BAL) lavages in non-human primates (NHP) for Molnupiravir, Paxlovid, and the combination of both drugs.^36^ Rhesus macaques were randomly separated into groups of five, where they received vehicle, Molnupiravir, Paxlovid, or a combination of both drugs. All animals were infected with the delta variant at a dose of 2 × 10^6^ TCID50 split between the nose and trachea and all treatments started 12 hours post-infection via an oral gavage. Five animals received a dose of 130 mg/kg of Molnupiravir every 12 hours, and another five received a dose of 20 mg/kg of Paloxivid. Finally, five animals received both 130 mg/kg of Molnupiravir and 20 mg/kg of Paxlovid simultaneously every 12 hours. In all cases, treatment consisted of two doses daily and continued until day 3.5 post-infection. Viral load observation for all monkeys was gathered via PCR and TCID50 for nasal and BAL as described in the treatment manuscript.^36^

### PK data

Pharmacokinetic data for Molnupiravir (EIDD-2801) was obtained from a study which assessed its efficacy in reducing viral load in non-human primates (NHPs) infected with influenza.^83^ A total of eight NHPs received a single 100 mg/kg dose, and plasma drug concentrations were monitored over the following 24 hours. Animals were monitored at 0.25, 0.5, 1,2,3,4,6,8,12,18 and 24 hours post dose.

Similarly, PK data for nirmatrelvir was sourced from,^75^ a study focusing on drug toxicity evaluation in NHPs. Plasma concentrations were measured for up to 15 days (which was the duration of the treatment) following twice-daily administration. Animals were monitored at 0.5, 1,2,4,6,7,12 and 24 hours post dose. Eighteen NHPs were divided into three dosage groups, with six receiving 20 mg/kg, six receiving 50 mg/kg, and six receiving 300 mg/kg.

### Nirmatrelvir model

We used a standard within-host framework to capture SARS-CoV-2 dynamics observed in both TCID50 and PCR data. The model consists of a system of ordinary differential equations (ODEs) with six main compartments: susceptible cells (*S*), exposed cells (*E*), infected cells (*I*), refractory cells (*R*), infectious virus (*V*_*pfu*_), non-infectious virus(*V*_*NI*_). Infected cells (*I*) die at a constant rate. These cells produce virions at a rate *p*, with a portion *μ* classified as infectious (*V*_*pfu*_) and the remainder as non-infectious (*V*_*NI*_). Nimatrelvir blocks viral production according to efficacy *ε*(*t*) which is a function of the joint PK and PD models.

To account for innate immune responses, we indirectly incorporated the effects of interferon secretion. When susceptible cells encounter infected cells, they may become refractory to infection at a rate *ϕIS*. However, this protection is temporary and wanes over time at a rate *ρR*.

In consequence, the model has the following form,

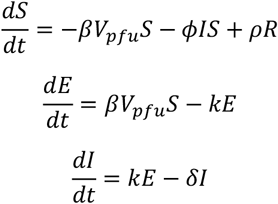

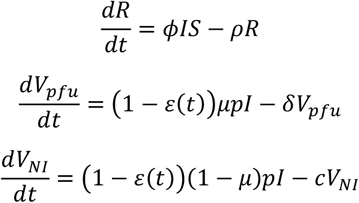

### Molnupiravir model

The molnupiravir infection model has equivalent infection assumptions and consists of a similar system of ordinary differential equations (ODEs) with the same main compartments: susceptible cells (*S*), exposed cells (*E*), infected cells (*I*), refractory cells (*R*), infectious virus (*V*_*pfu*_) and non-infectious virus(*V*_*NI*_). Molnupiravir makes infected cells produce mutated / non-infectious virus (*V*_*mut*_) according to drug efficacy *ε*(*t*) which is dependent on the PKPD model. The remaining virus is either *V*_*NI*_ or *V*_*pfu*_.

Thus, the model follows the structure:

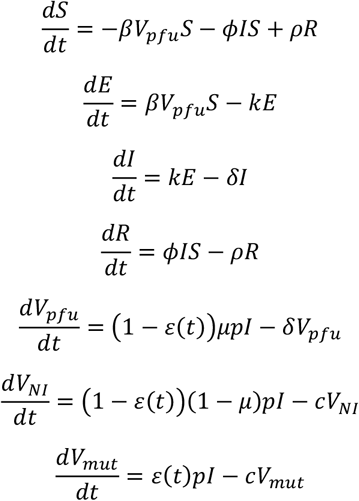

### PK models

For both type of drugs, we used a two-compartmental model PK which includes the amount of drug in the gastrointestinal tract (*A*_*GI*_), the plasma compartment (*A*_*P*_) and lung tissue (*A*_*L*_). The drug was administrated orally using a french catheter directly to the GI tract, where it is assumed to get absorbed into the blood at a rate *k*_*a*_. Once in the blood, the drug will transfer to tissue at a rate *k*_12_. As the drug is metabolized it comes back to the plasma from the lungs at a rate *k*_21_, where it is eliminated from the body at the rate *k*_*cl*_*k*. The model in the form of ODEs is the following:

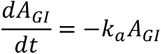

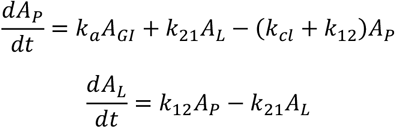

We used Monolix and a mixed-effect population approach with their respective PK data from each drug, with the same initial conditions of *A*_*GI*_ = *Dose, A*_*P*_ = 0, *A*_*L*_ = 0. We fitted the 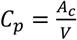 to the plasma concentration data, where V is the estimated plasma volume.

### PD models

The data on drug efficacy we used the parameters that were obtained from experiments performed in prior published work.^49,71^ For Paxlovid data the parameters were retrieved from^49^ and for Molnupiravir parameters were obtained from^71^. We used the parameters obtained from an in vitro efficacy data collected at different doses of nirmatrelvir. The *E*_*max*_, the invitro *EC*_50_ and the hill value *n* were fixed based on the values estimated in the experiment performed to evaluate the efficacy of nirmatrelvir in Calu-3 cells infected with the delta variant. The invivo *EC*_50_ was estimated based on the potency reduction factor defined as 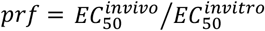. The Emax, Hill coefficient (n), and IC50 values derived from human Calu-3 cells are considered pharmacodynamically relevant for non-human primate (NHP) studies, as these parameters reflect the intrinsic activity of the compound at a conserved biological target. Calu-3 cells, which model human airway epithelium, express key host factors and cellular pathways that are functionally similar in NHPs. Given the high degree of conservation in drug target engagement and downstream antiviral mechanisms across species, in vitro PD parameters from human cell-based assays can serve as reliable surrogates for in vivo PD modeling in NHPs. This translational approach is particularly justified when the mechanism of action is host-directed and less susceptible to species-specific differences.

### Nonlinear mixed effects model and model fitting

To estimate the model parameters and fit the models to the PCR and TCID50 observations simultaneously, we used a non-linear mixed effects approach. Under this approach, a PCR or TCID50 observation from a ferret *i* at a time *j* is modeled as log_10_*Y*_*ij*_ = log_10_ *f*_*PCR*_(*t*_*ij*_, *θ*_*i*_) + *∈*_*PCR*_ and log_10_ *Z*_*ij*_ = *f*_*PFU*_(*t*_*ij*_, *θ*_*i*_) + *∈*_*PFU*_. Here, *f*_*PCR*_ and *f*_*TCID*50_ are defined as *f*_*PCR*_ = *V*_*ni*_(*t*_*ij*_, *θ*_*i*_) + *V*_*inf*_(*t*_*ij*_, *θ*_*i*_) and *f*_*PFU*_ = *V*_*inf*_(*t*_*ij*_, *θ*_*i*_) based on the solution for the ODE model variables that describe non-infectious *V*_*ni*_ and infectious virus *V*_*inf*_. Meanwhile, *θ*_*i*_ is a vector that contains the parameters values for ferret *i*, and *∈*_*PCR*_ and *∈*_*PFU*_ are the measurement errors for the log_10_-transformation of the PCR and TCID50 data. We assumed *θ*_*i*_ comes from a probability distribution with median or fixed effects *θ*^*pop*^ and random effects *η*_*i*_∼*N*(0, *σ*_*θ*_). All modeled parameters are log-normally distributed in the population, i.e., *θ*_*i*_ = *θ*^*pop*^*e*^*η_ij_*^, except β and *p* that were normally (log_10_ *θ_i_* = *θ*^*pop*^ + *η_i_*) and δ logit-normally 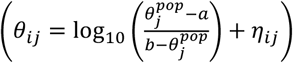 distributed, respectively.^84-87^

We used the Stochastic Approximation Expectation Maximization (SAEM) algorithm which is set in the Monolix software to estimate the population and error parameters of our model, and a Markov Chain Monte Carlo approach to estimate the individual parameter values based on the population parameters distribution. In this model, we assume the value of *t* = 0 as the time of infection intranasally on the NHP with initial conditions *V*_*inf*_(0) = 0, *V*_*NI*_ (0) = *V*_*zero*_. Since we are studying the infection dynamics in NHP, we assumed *S*(0) = 2.6*x*10^6^ cells/ml, based on an estimated percentage of cells that express the sufficient receptor for the virus to fuse and enter the cell based on a human infection model.^88^ Finally, we ran the SAEM algorithm 10 times (assessments) for each model where we randomly selected the initial value of the estimated parameters. From the 10 simulations of the SAEM algorithm we calculated the log-likelihood (log_10_ Γ) to evaluate the goodness of fit. The assessment with the highest log Γ was used to compute the value of the Akaike Information Criteria (AIC) which was obtained by the following formula *AIC* = −2 max(log_10_ Γ) + 2*m*, where *m* is the number of parameters estimated. In the electronic supplementary material, **table S1** contains the equations for all the models evaluated, with their respective AIC and log Γ values.

